# Structural basis for the assembly of the DNA polymerase holoenzyme from a monkeypox virus variant

**DOI:** 10.1101/2022.12.16.520684

**Authors:** Yaning Li, Yaping Shen, Ziwei Hu, Renhong Yan

**Affiliations:** Center for Infectious Disease Research, Westlake Laboratory of Life Sciences and Biomedicine, Key Laboratory of Structural Biology of Zhejiang Province, School of Life Sciences, Westlake University, Hangzhou 310024, Zhejiang Province, China; Department of Biochemistry, Key University Laboratory of Metabolism and Health of Guangdong, School of Medicine, Southern University of Science and Technology, Shenzhen 518055, Guangdong Province, China; Beijing Advanced Innovation Center for Structural Biology, Tsinghua-Peking Joint Center for Life Sciences, School of Life Sciences, Tsinghua University, Beijing 100084, China

## Abstract

The ongoing pandemic caused by a monkeypox virus (MPXV) variant has spread all over the world and raised great public health concerns. The DNA polymerase F8 of MPXV, associated with its processivity factors A22 and E4, is responsible for viral genome replication in the perinuclear sites of the infected cells as well as a critical target for developing antiviral drugs. However, the assembly and working mechanism for the DNA polymerase holoenzyme of MPXV remains elusive. Here, we present the cryo-EM structure of the DNA polymerase holoenzyme F8/A22/E4 from the 2022 West African strain at an overall resolution of 3.5 Å and revealed the precise spatial arrangement. Surprisingly, unlike any other previously reported B-family DNA polymerase, the holoenzyme complex is assembled as a dimer of heterotrimers, of which the extra interface between the thumb domain of F8 and A22 shows a clash between A22 and substrate DNA, suggesting an auto-inhibition state. Supplying an exogenous double-stranded DNA could notably shift the hexameric form into a trimeric form, which exposes the DNA binding site of thumb domain and might represent a more active state. The structures provide a molecular basis for the design of new antiviral therapeutics that target the MPXV DNA polymerase holoenzyme.

## Introduction

A monkeypox virus (MPXV) variant outbreak has rapidly spread over 110 countries and confirmed more than 80 thousand cases (as of Nov. 26^th^, 2022) since early May of 2022 (https://www.cdc.gov/poxvirus/monkeypox/response/2022/world-map.html) (*1–3*). The World Health Organization (WHO) has declared that the global monkeypox pandemic represents a public health emergency of international concern, calling for research on vaccines, therapeutics, and other tools against this virus (*4*).

MPXV is a member of *Poxviridae*, a family of enveloped, double-stranded DNA (dsDNA) viruses that also includes the variola virus, the causative agent of smallpox that has killed millions of people in the 20^th^ century (*5–7*). Phylogenetic analyses show the West Africa (WA) and the Congo Basin (CB) Clades (*8, 9*). The fatality rates for the CB clade are typically in the range of 1% to 5% whereas the patients suffered from WA clade cases are rarely fatal in non-immunocompromised patients (*10*). Genome sequencing identified the 2022 outbreak of MPXV diverged from a 2018 WA strain, but contains about 50 extra mutations, implying the accelerated evolution of MPXV (*11*).

MPXV is a kind of contagious zoonotic virus that was first isolated from monkeys and could be transmitted incidentally to humans when in close contact with infected animals (*8, 12, 13*). MPXV could also infect many kinds of mammals including the rope squirrels, Gambian pouched rats, African hedgehogs, chimpanzees, and so on (*9, 14*). Human-to-human spread often involves direct contact with infectious rash, sores, and scabs (*15, 16*). Besides, many confirmed cases in the current MPXV outbreak are traced to sexual transmission (*17, 18*).

During productive infection, the *Orthopoxvirus*, such as MPXV, replicates its genomes in the perinuclear sites of the infected cells using a set of virus-encoded proteins (*19–21*). Similar to vaccinia virus (VACV), another well-studied member of *Poxviridae*, the DNA polymerase holoenzyme of MPXV encoded by the viral genome contains three proteins: the uracil-DNA glycosylase E4, the processivity factor A22, and the DNA polymerase F8 belonging to the B-family polymerase, which are corresponding to D4, A20, and E9 in VACV, respectively (*22–24*). A22 works as a bridge for connecting F8 and E4, among which E4 is crucial for the association of viral DNA templates with enzymes to process efficiently (*25–29*).

Many attempts targeted to the DNA polymerase holoenzyme of *Poxviridae* are developed, such as the acyclic cytosine phosphonate analogue: cidofovir (CDV) and the cidofovir derived prodrug brincidofovir (also known as CMX001), but DNA polymerase mutant variant conferred resistance to these drugs (*30–33*). There are 6 extra mutations (L108F, W411L, T428I, S484A, V501I, and D785N) located on F8 DNA polymerase from 2022 WA strain of MPXV compared with previous strains, raising a greater risk for drug resistance (Fig.1A) (*11, 34*). Notably, the expression of DNA polymerase E9 of VACV is strictly controlled during infection process (*35, 36*). The levels of E9 mRNA transcription and translation peaked at about 2 to 3.5 hours after infection and then decline, becoming barely detectable by 5 to 6.5 hours post-infection, probably for transition into late gene expression (*35*). However, whether the DNA polymerase activity of E9 of VACV or F8 of MPXV is simultaneously regulated during the infection process remains unclear.

**Fig. 1.**
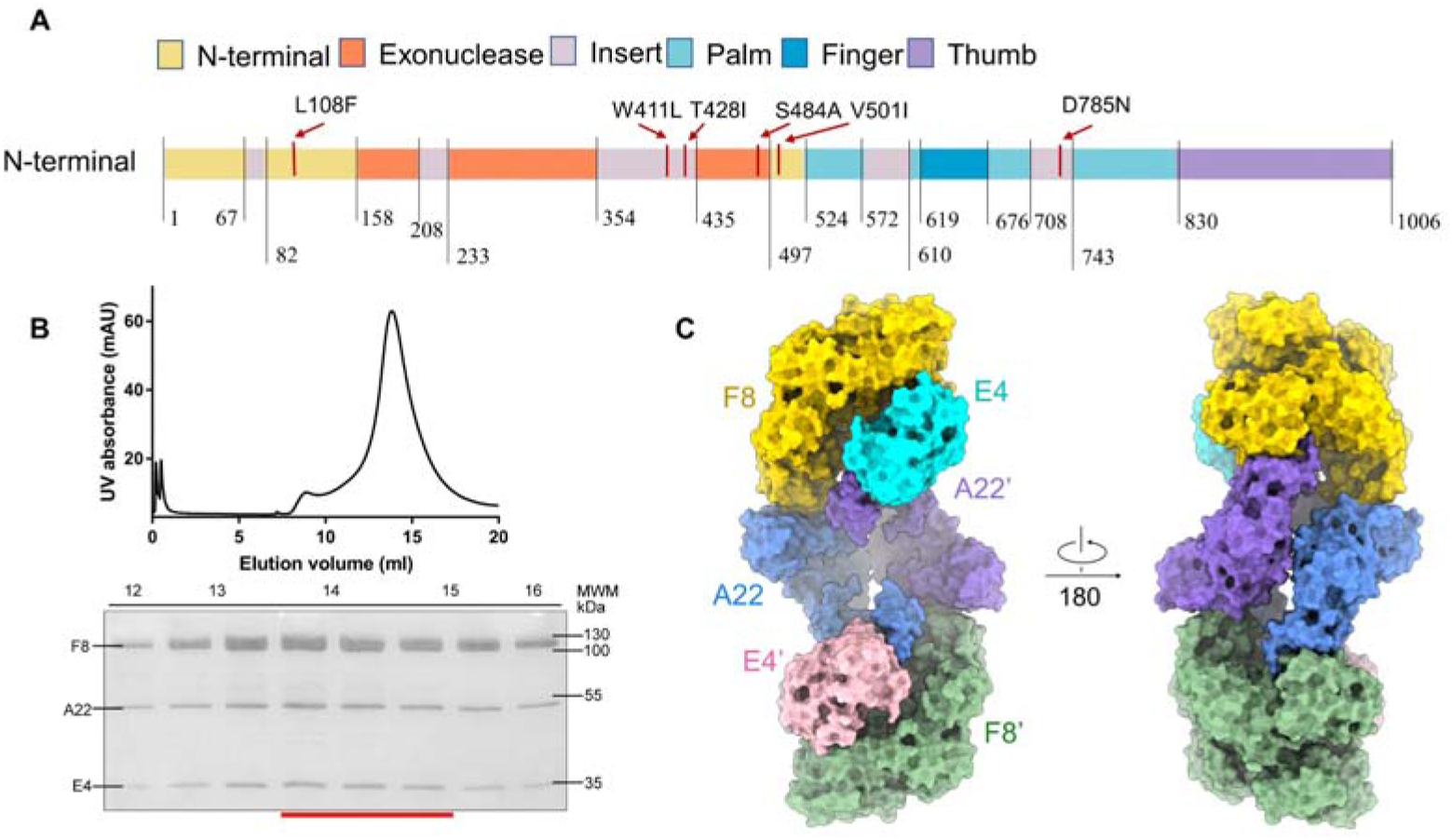
Biochemical characteristics of DNA replication machine of MPXV. (A) Domain organization of F8L. Mutations in 2022 WA strain are marked by red lines. (B) Representative SEC purification of the F8-A22-E4 complex. SDS-PAGE was visualized by Coomassie blue staining and fractions for cryo-EM analysis were marked by red line. (C) Overall surface presentation of domain-colored cryo-EM structures of DNA replication machine of monkeypox. F8, A22, and E4 in protomer A are colored gold, blue, and cyan respectively, and the other protomer is colored green, purple, and pink.

Despite the insights gained from VACV structural research, including the crystal structures of DNA polymerase E9 alone, the processivity factor A20(1-50 a.a)/D4 complex, C-terminal Domain of A20 (304-426 a.a), and low-resolution model of E9/A20/D4 (*23, 25, 28, 29*), there remains very limited knowledge concerning the precise assembly and action mechanism for the holoenzyme of *Poxviridae*. Here, we present the cryo-EM structure of the DNA replication holoenzyme F8/A22/E4 from the 2022 WA strain (Genbank ID: ON563414.3) at an overall resolution of 3.5 Å. Surprisingly, unlike any other previously reported B-family polymerase, the holoenzyme complex is assembled as a dimer of heterotrimers. To the best of our knowledge, this is the first time that the hexameric assembly of B-family polymerase has been revealed. Structural analysis illustrated that in the hexameric form of the DNA polymerase holoenzyme from MPXV, the DNA polymerase F8 directly interacts with the uracil-DNA glycosylase E4 in contrast to the previous low-resolution models from VACV, which assumed the complex to be elongated in shape and the DNA polymerase and uracil-DNA glycosylase are separated (*26*). The extra interface between the thumb domain of F8 and the middle Neck domain of A22’ might preclude the binding of DNA substrate, this could hint at an inactive conformation. When supplied with an exogenous double-stranded DNA (dsDNA), the hexameric form of the holoenzyme changed into a trimeric form notably, exposing the DNA binding site of thumb domain. Besides, the high-resolution structure of F8 enables detailed analysis of each domain and newly emerged mutations, which further provides a good framework for estimating drug availability.

## Structural determination of F8/A22/E4 holoenzyme

The full-length open reading frames of F8L/A22R/E4R from the 2022 WA strain were co-expressed in HEK293F cells and purified through tandem affinity resin and size exclusion chromatography. The complex was eluted in a single mono-disperse peak, indicating high homogeneity (Fig. 1B). Details of cryo-sample preparation, data acquisition, and structural determination are provided in the Materials and Methods and sections of supplementary materials (Figs. S1-S5 and Table S1). A 3D reconstruction was obtained at an overall resolution of 3.5 Å from 391,085 selected particles. Thus immediately revealed the dimer of heterotrimers’ architecture (Fig. 1C).

The high resolution structure supported reliable model building. For F8 polymerase, the 934 side chains out of 1006 residues could be assigned, which contain the N-terminal, exonuclease, palm, finger, thumb domains and five “poxvirus-specific” inserts (Insert 0 to Insert 4) (Fig. 1A and 2A) (*26*). The newly emerged mutations are clearly mapped on the structure (Fig. S6A). The overall architecture of F8 shows an open conformation and is similar to that of VACV with a root mean square deviation (RMSD) value of 3.94 Å for 934 Cα, of which the thumb domain is significantly shifted due to the interaction of the Neck domain of A22’ (Fig. 2D) (*23*). The full-length structure of A22 is firstly resolved and connected to F8 and E4. Besides, the overall structure of E4 is also similar to its homologous protein, D4, in VACV (*26*).

**Fig. 2.**
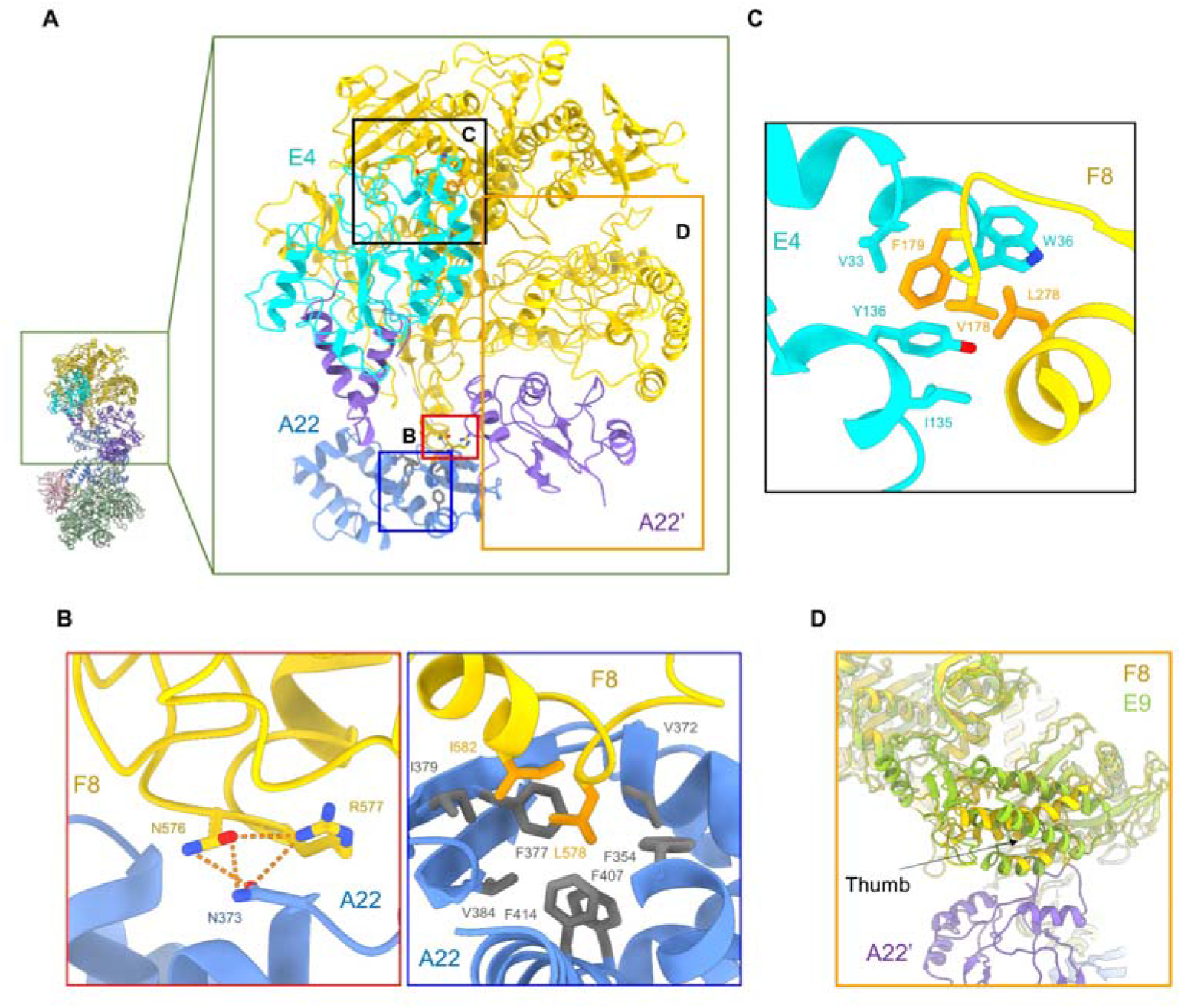
The multiple interaction interfaces in F8-A22-E4 hexameric complex. (A) Overall structure of F8-A22-E4 complex. Interaction interfaces between F8 and A22 in one protomer are lined out by red and blue boxes, which are shown in (B); The interacted residues in F8 and E4 are lined out by black box and showed in (C); Structural comparison with E9 DNA polymerase from vaccinia virus (PDB ID: 5N2E) shows the thumb domain is regulated by the interaction of A22’. F8, E4, A22, and A22’ are colored gold, cyan, blue and purple, respectively. E9 is colored yellow green.

## Multiple interfaces of holoenzyme

The A22 could be divided into three parts: the N-terminal domain, the middle Neck domain, and the C-terminal domain. The linker (46-101 a. a) between N-terminal domain and Neck domain is invisible when high resolution is reached, but the low-resolution threshold could help identify the direction of this chain (Fig. S6B).

For clarity, two protomers are referred to as protomer A: F8/A22/E4 and protomer B: F8’/A22’/E4’. The overall architecture exhibits as a central symmetry, creating an extra interface: the Neck domain of A22 and thumb domain of F8’ (Fig. 2). Because of flexibility, the density of the interface between Neck domain of A22’ and thumb domain of F8 is not good enough to support detailed model building (Fig. 2D and Fig. S4D). Overall, A22 plays a bridge role in binding the N-terminal domain of A22 to E4’ and C-terminal domain of A22 interaction with the poxvirus specific Insert 3 region of F8.

Since the overall architecture exhibiting as a central symmetry, the focused refinement is applied on protomer A, which improves the resolution up to 3.3 Å, enabling detailed analysis (Fig. 2). The interface between F8 and C-terminal domain of A22 is mainly mediated by the short helices of poxvirus specific insert 3 region that inserts into the hydrophobic pocket of A22. The extensive hydrophobic network contains the Leu578 and Ile582 of F8 and Phe354, Val372, Phe377, Ile379, Val384, Phe407, and Phe414 of A22. Meanwhile, Asn576 and Arg577 of F8 are respectively hydrogen bonded (H-bonded) with the Asn373 of A22 (Fig. 2B). Additionally, the interface between F8 and E4 is mainly mediated by the hydrophobic interactions involved in the Val178, Phe179, and Leu278 of F8 and Val33, Trp36, Ile135, and Tyr136 of E4 (Fig. 2C). Compared with E9, the homologue polymerase in VACV, the thumb domain is significantly shifted probably due to the interaction with the Neck domain of A22’ (Fig. 2D).

## Distinct organization pattern

Previous sequence analysis of the VACV polymerase has revealed the homology to the replicative B-family of PALM polymerases (Pol α and Pol δ) encoded by mammalian cells and a variety of DNA viruses such as Herpes simplex virus (HSV) and bacteriophage RB69 (*23, 37*). The B-family polymerases usually accomodate some co-factors for processivity (*38*). However, the organization pattern of different members of B-family polymerases varies widely (Fig. 3A-D). The bacteriophage RB69 acquires the trimeric sliding clamp factor PCNA through the C-terminal thumb domain of polymerase (Fig. 3B) (*39*). Similarly, the PCNA-related processivity factor (UL42) of HSV also binds to the C-terminal thumb domain of the polymerase (UL30) (Fig. 3C) (*40*). While in eukaryotic polymerase δ, which acquires more processivity factors, such as Pol31 and Pol32, involved in the replication process, retains the ability to bind PCNA through the elongated C-terminal domain (Fig. 3D) (*41*). Compared with these B-family polymerases, the typical features for the replication machine in MPXV are involved with the base-excision enzyme, the uracil-DNA glycosylase E4 and the dimer of heterotrimeric organization pattern (Fig. 3A-D).

**Fig. 3.**
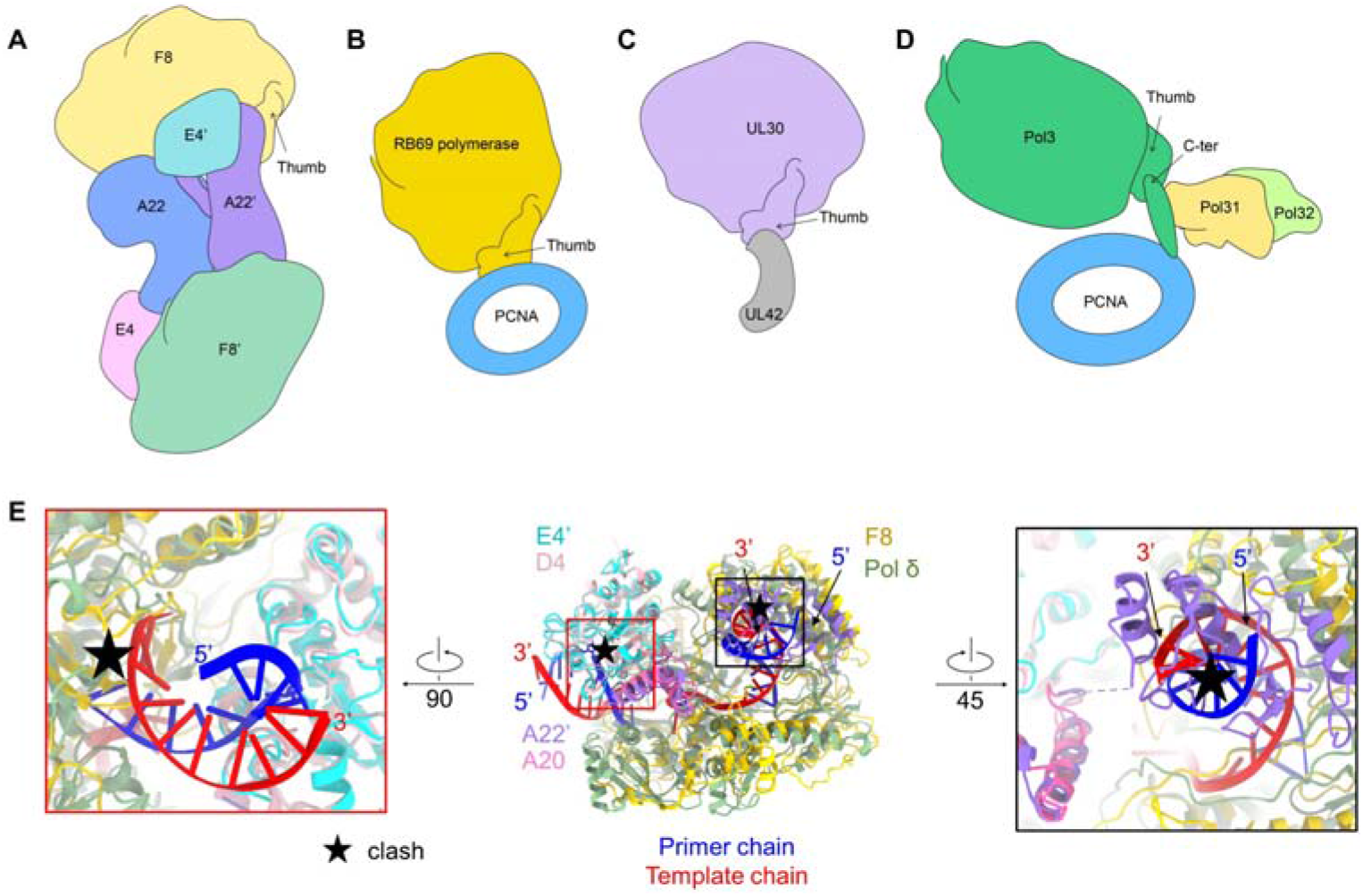
The models of different DNA replication machines. (A) Model of DNA polymerase holoenzyme from MPXV. (B) Archaeal and phage DNA polymerase. The model was drawn according to polymerase structure (PDB ID: 1CLQ) and PCNA structure (PDB ID: 1B8H). (C) HSV DNA polymerase. The model was drawn according to UL42 structure (PDB ID: 1DML) and UL30 structure (PDB ID: 2GV9). (D) S. cerevisiae polymerase. The model was drawn according to a complex structure (PDB ID:7KC0). (E) Structural comparison with DNA polymerase delta of yeast (PDB ID: 3IAY) and D4-A20 complex of vaccinia virus (PDB ID: 4YIG) shows there would be a clash between DNA and F8 or A22’. F8, A22, A22’, and E4 are colored gold, blue, purple, and cyan, respectively. D4 and A20 of vaccinia virus are colored pink and magenta. DNA polymerase delta of yeast is colored dark sea green. The template and primer chains of DNA are colored red and blue.

The unexpected organization pattern of the DNA polymerase holoenzyme of MPXV raises the concern of the working mechanism. In VACV, the uracil glycosylase activity of E4 has been shown to be active in the context of the polymerase holoenzyme and plays a role like the sliding clamp, as evidenced by the *in vitro* replication assay that could generate abasic sites in a uracil-containing oligonucleotide and viral DNA replication and repair could be coupled (*42*). However, there is an obvious clash between the substrate uracil-containing DNA and F8 in the solved holoenzyme structure of MPXV, suggesting an inactive form of E4 in this structure (Fig. 3E, left panel). To be noticed, when aligning the F8 with yeast DNA polymerase δ, the thumb domain of F8 is engaged by the Neck domain of A22’ and also shows a clash between A22’ and substrate dsDNA, suggesting an inhibitory effect of A22 on the elongation of DNA polymerase in this structure (Fig. 3E, right panel). Taken together, these findings suggest the hexameric form of DNA polymerase holoenzyme from MPXV might represent as an inactive conformation.

## Two different states of holoenzyme reveal the underlying working mechanism

To explore the molecular mechanism of DNA replication by the MPXV, we tried to incubate the dsDNA substrate (See Materials and Methods section) with the hexameric complex form. To our surprise, the addition of the exogenous dsDNA leads to the increase of a latter peak on gel filtration and it still contains the bands of F8/A22/E4 on the SDS-PAGE, which might represent another state of the holoenzyme (Fig. 4A). We then collected the latter peak fractions and prepared the cryo-EM sample. Structural determination reveals the structure of ternary form with F8/A22/E4 at the resolution of 3.1 Å (Fig. 4B and Figs. S3, S5 and Tabls S1). However, both peak fractions could not get the DNA substrate bound state.

**Fig. 4.**
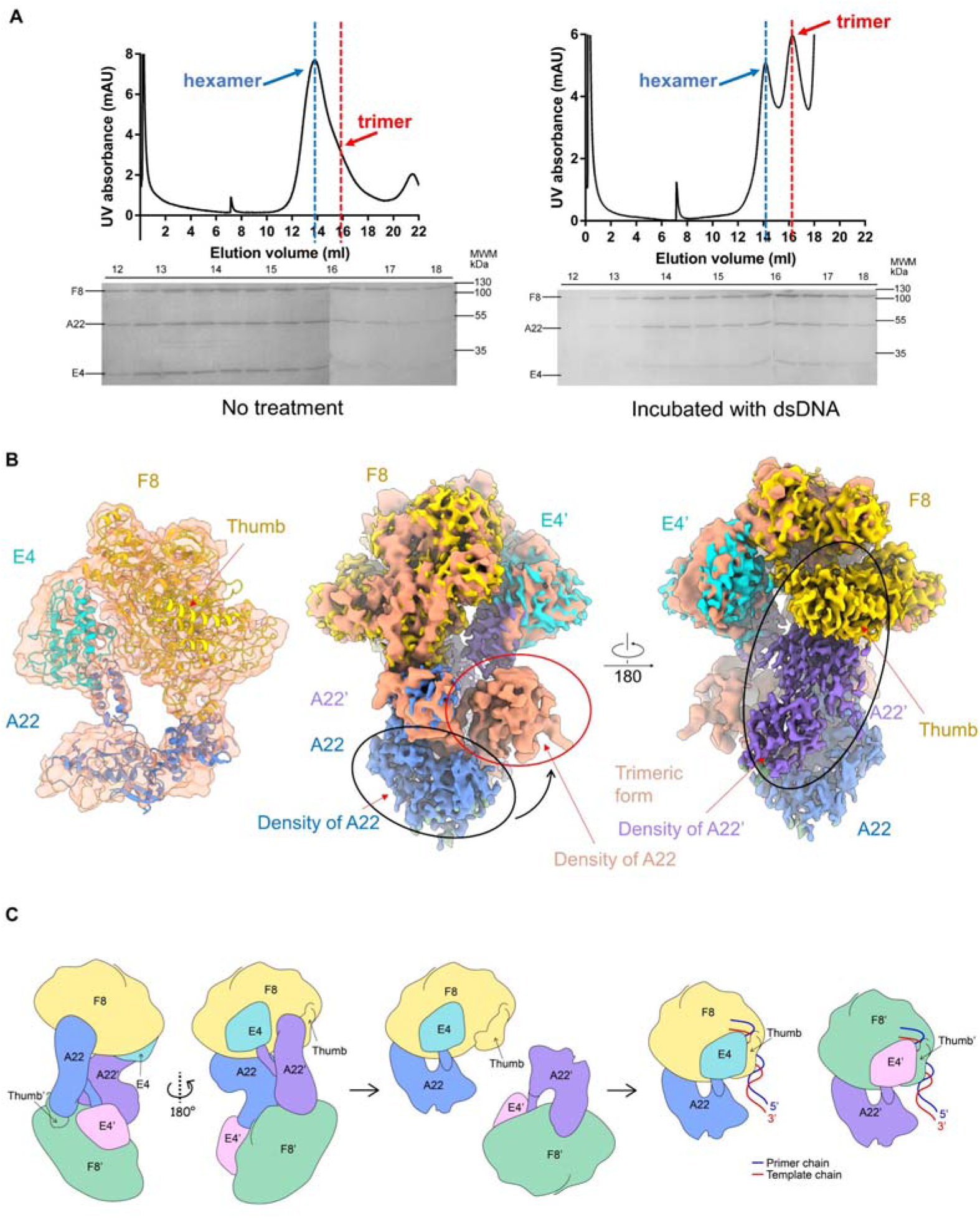
The putative working model of F8-A22-E4 complex. (A) The oligomer state shift of F8-A22-E4 complex from hexamer to trimer can be induced by incubation with dsDNA. The peak of hexamer and trimer are marked by blue line and red line respectively on SEC analysis. The major peak of F8-A22-E4 complex shifts backward when incubated with dsDNA. (B) *Left panel:* Map and model of trimeric form complex. The map of trimeric form complex is colored light salmon. The model styled cartoon and F8, A22, and E4 are colored gold, blue, and cyan, respectively. *Middle and right panel:* Map comparison of trimeric form and hexameric form complex shows A22 undergoes dramatically conformational change during oligomer state shift. (C) A proposed model for auto-inhibition, activation and the DNA replication mechanism catalyzed by F8-A22-E4 complex.

The ternary structure of F8/A22/E4 complex might exhibit as a more active state due to the loss of the interface between the thumb domain of F8 and A22’, thus releasing the dsDNA binding site of F8 (Fig. 4B). The transition from hexameric form to trimeric form requires a dramatic movement of the N-terminal domain and Neck domain of A22, in which the dissociation of A22’ could lead to the replacement of the interface of A22’-E4 into A22-E4 (Fig. 4B and S6B).

## Discussion

The replication process of poxvirus is strictly controlled during the infection process. The peak of mRNA and protein levels of E9 in VACV is 2 to 3.5 hours after infection and declines to be undetectable by 5 to 6.5 hours, aiming for transition into late gene expression state (*35*). Besides, BAF (barrier to autointegration factor) dimers can bind tightly to dsDNA and occlude transcription and replication of VACV, however, B1-mediated phosphorylation of BAF prevents this inhibitory effect (*43, 44*). Many studies have shown the polymerase alone of poxvirus is intrinsically distributive but the processivity factor could render it highly processive, suggesting the critical role of A22 and E4 (*22, 45*). Whether the DNA replication process of poxvirus is timely regulated remains elusive. Here, we found the two different states of DNA replication holoenzyme from MPXV, that could be induced by the dsDNA to transition an inactive hexameric form into a more active trimeric form. We proposed a possible working model for the DNA replication holoenzyme by the F8/A22/E4 complex based on our findings (Fig. 4C). In the proposed model, the auto-inhibitory hexameric form of holoenzyme could not exert the replication function since the DNA binding site of F8 and E4 are not exposed. Once the dsDNA substrate or another factor binds to the holoenzyme, it could trigger the conformational change into a ternary complex, releasing the DNA binding site of thumb domain, then a replication process happened that change the open conformation of F8 into closed conformation to bind the template-primer strands and E4 might also experience a conformational change to capture DNA.

We also wonder about the role of CDV and CDV derived prodrug CMX001 against MPXV (*46, 47*). It’s reported that CDV could be metabolized into the triphosphate form (cidofovir diphosphate, CDVpp) to be served as competitive inhibitor of deoxycytidine triphosphate (dCTP) binding to inhibit a growing DNA strand and destabilize the dsDNA of virus (*33, 48, 49*). The previously reported structures of B-family polymerases in complex with DNA substrate, with or without an incoming nucleotide mimicking elongation mode, indicating that large domain movements happen upon DNA binding leading to a closure of the structure compared to the open conformation (*50*). To study the effect of CDV on viral DNA polymerase, molecular docking analysis was carried out to estimate the CDVpp binding between E9 of VACV and F8 of MPXV. We used the yeast polymerase δ structure with bound template and complementary DNA (PDB ID: 3IAY) to generate the elongation mode of F8 and E9 by SWISS-MODEL (Fig. S7) (*50, 51*). From our docking results, CDVpp exhibited considerable binding affinity to closed conformation of E9 and F8, −7.3 Kcal/mol and −7.6 Kcal/mol, respectively. We found that CDVpp is incorporated into the DNA chain opposite the dG in a state strikingly similar to incoming dCTP. CDVpp enters into the active cavity of F8 or E9 via conventional H-bonds to main chain of Ser552, Leu553, and side chain of Lys661, which are also critical to stabilize the incoming natural dCTP (Fig. 5A). These results suggested that CDV and its analogue have high effectiveness in the treatment of MPXV infection. Another dCTP competitor, the nucleotide analogue 1-beta-D arabinofuranosylcytosine (AraC) also shows a reasonable binding affinity with E9 (−6.4 Kal/mol) and E8 (−6.7 Kal/mol), indicating a putative therapeutic candidate (*52*). AraC is capable of acting as direct chain terminators, which involves in the hydrogen bonding network with main chain of Leu553, Tyr554, and side chain of Asn665 located at the vicinty of the dCTP-binding pocket (Fig. 5B).

**Fig. 5.**
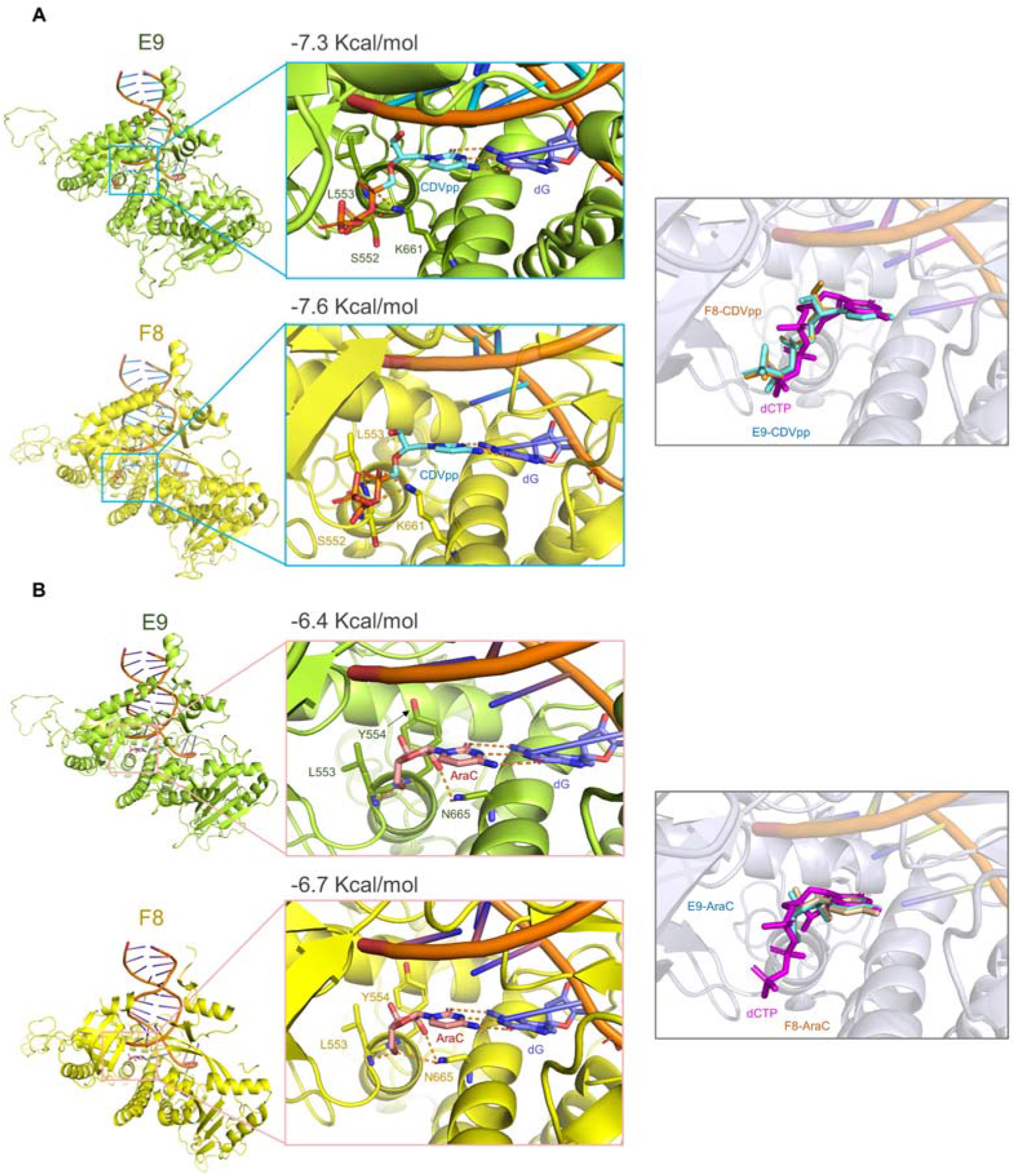
Analysis of drug inhibitors on E9 and F8. (A) Molecular docking of CDVpp binding mode in E9 and F8, respectively. The interaction residues are shown as sticks. E9 and F8 are colored green and yellow, respectively. The structure on the right is the comparative analysis of dCTP (purple) and CDVpp (blue) in the E9 and F8, respectively. (B) Molecular docking of AraC in E9 and E8, respectively. The interaction residues are shown as sticks. The alignment analysis on the right is also the comparative analysis of dCTP (purple) and AraC in the E9 and F8, respectively.

The remodeling structure of F8 and our newly reported structures provide an important clue into the putative effect of emerged mutations of MPXV (Fig. S8). For an overall view, L108F mutation is close to the template chain of dsDNA and may strengthen the hydrophobic interaction with base in template chain. W411L and T428I are both located on the insert 2 region of F8, which may interact with the finger domain upon dsDNA binding as does ‘polymerase associated domain’ (PAD) in Y-family polymerases and probably serve as a regulatory factor binding site (*34, 53*). The surface exposed hydrophobic W411L and T428I might affect the binding affinity of putative regulatory factor.

In sum, the high-resolution structure of the MPXV DNA polymerase holoenzyme reveals the mode of processivity factor binding and provides an understanding of putative working mechanisms, although the dynamic conformational change of the holoenzyme still requires further structural and functional research in order to obtain more functional states. The identification of the interfaces of F8/E4 and F8/A22 might facilitate the design of new antiviral drugs that disrupt these interactions.

## Acknowledgments

We thank the Cryo-EM Facility of Southern University of Science and Technology (SUSTech) for providing the facility support. We thank Shuman Xu, Peiyao Li and Lutang Fu at the Cryo-EM Center of SUSTech for technical support in electron microscopy data acquisition. We thank the Cryo-EM Facility and Supercomputer Center of Westlake University for providing cryo-EM and computation support, respectively. We thank Zhenyuan Liu for technical support on computing environment. This work was funded by the Science, Technology and Innovation Commission of Shenzhen Municipality (JSGG20220226085550001 to R.Y.), and the National Natural Science Foundation of China (82202517 to R.Y.).

## Author contributions

R.Y. conceived the project. R.Y., Y.L., and Y.S. designed the experiments. Y.L. did the molecular cloning, protein purification, Cryo-EM data collection and processing, model building and DNA induced assay. Y.S. did the protein purification, Cryo-EM data collection and DNA induced assay. Z.H. did the molecular docking assay. All authors contributed to data analysis. R.Y. wrote the manuscript.

## Data and materials availability

Atomic coordinates and cryo-EM density maps of F8-A22-E4 in hexameric form (PDB: XXXX whole map: EMD-XXXXX, map focused on the protomer A: EMD-XXXX) and Atomic coordinates and cryo-EM density maps of F8-A22-E4 in trimeric form (PDB: XXXX whole map: EMD-XXXXX) have been deposited to the Protein Data Bank (http://www.rcsb.org) and the Electron Microscopy Data Bank (https://www.ebi.ac.uk/pdbe/emdb/), respectively. Correspondence and requests for materials should be addressed to R.Y.

## Supplementary Materials

Materials and Methods

Figures S1-S8

Table S1

Supplementary references (1-16)

