## Supplementary_Information for "Structural basis for the assembly of the DNA polymerase holoenzyme from a monkeypox virus variant"

### Materials and methods

#### Protein expression and purification

The full length F8 of Monkeypox virus (MPXV 2022 West African strain, GenBank: ON563414.3) was cloned into the pCAG vector (Invitrogen) with a C-terminal 10×His tag. The full length E4 and A22 (MPXV 2022 West African strain, GenBank: ON563414.3) were cloned into the pCAG vector (Invitrogen) with a N-terminal FLAG tag. All the plasmids used to transfect cells were prepared by GoldHi EndoFree Plasmid Maxi Kit (CWBIO).

The recombinant protein was overexpressed using the HEK293F mammalian cells at 37℃ under 5% $CO_2$ in a Multitron-Pro shaker (Infors, 130 rpm). When the cell density reached 2.0 $\times 10^6$ cells/mL, the plasmid was transiently transfected into the cells. To transfect one liter of cell culture, about 1.5 mg of the plasmid was premixed with 3 mg of polyethylenimines (PEIs) (Polysciences) in 50 mL of fresh medium for 15 mins before adding to cell culture. Cells were collected by centrifugation at 4000×g for 10 mins after sixty hours transfection and resuspended in a buffer containing 25 mM HEPES (pH 7.5), 150 mM NaCl, and mixture of three protease inhibitors, aprotinin (1.3 μg/ml, AMRESCO), pepstatin (0.7 μg/ml, AMRESCO) and leupeptin (5 μg/ml, AMRESCO).

For F8/A22/E4 complex purification, cells were lysed by sonication and cell debris was removed by centrifugation at 18700×g for 45 mins. The supernatant was loaded onto anti-FLAG M2 affinity resin (Sigma). The resin was washed with the wash buffer containing 25 mM HEPES (pH 7.5), 150 mM NaCl, following by protein eluted with wash buffer plus 0.2 mg/mL FLAG peptide. Then elution of anti-FLAG M2 affinity resin was further purified with Ni-NTA affinity resin (Qiagen). Wash buffer and elution buffer of nickel resin was wash buffer mentioned above plus 10 mM and 300 mM imidazole respectively. Then the protein complex was subjected to size-exclusion chromatography (Superose 6 Increase 10/300 GL, GE Healthcare) in buffer containing 25 mM HEPES (pH 7.5), 150 mM NaCl. The peak fractions were collected and concentrated for EM analysis.

**DNA induced conformational change assay**

DNA duplex used for assay are the 12/16 primer template from Pol δ-DNA complex (PDB ID:3IAY), containing the 12-nt oligonucleotide primer (5′-ATCCTCCCCTAC-3′) annealed to the 16-nt template (5′-TAAGGTAGGGGAGGAT-3′)(*1*).

Fresh F8/A22/E4 complex from peak fractions of size-exclusion chromatography are incubated with the DNA duplex at a molar ratio of about 1:3 plus 2 mM $MgCl_2$ for one hour. Protein incubated with 2 mM $MgCl_2$ and same volume buffer just without DNA duplex is used as negative control. Then the mixture was subjected to size-exclusion chromatography (Superose 6 Increase 10/300 GL, GE Healthcare) in buffer containing 25 mM HEPES (pH 7.5), 150 mM NaCl.

**Cryo-EM sample preparation and data acquisition**

For complex in hexameric form, the fresh protein complex was concentrated to ~3 mg/mL and applied to the grids. For complex in trimeric form, the latter peak from DNA induced conformational change assay was collected and concentrated to ~3 mg/mL. Aliquots (3.3 μL) of the protein were placed on glow-discharged holey carbon grids (Quantifoil Au R1.2/1.3). The grids were blotted for 3.0 s or 3.5 s and flash-frozen in liquid ethane cooled by liquid nitrogen with Vitrobot (Mark IV, Thermo Fisher Scientific). The prepared grids were transferred to a Titan Krios operating at 300 kV equipped with Gatan K3 detector and GIF Quantum energy filter. Movie stacks were automatically collected using AutoEMation(*2*), with a slit width of 20 eV on the energy filter and a defocus range from -1.4 µm to -1.8 µm in super-resolution mode at a nominal magnification of 81,000×. Each stack was exposed for 2.56 s with an exposure time of 0.08 s per frame, resulting in a total of 32 frames per stack. The total dose rate was approximately 50 $e^-/Å^2$ for each stack. The stacks were motion corrected with MotionCor2[ref.](*3*) and binned 2-fold, resulting in a pixel size of 1.095 Å/pixel or 1.072 Å/pixel. Meanwhile, dose weighting was performed(*4*). The defocus values were estimated with Gctf(*5*).

**Data processing**

The Cryo-EM structure of the complex was solved in Relion 3.0.6 and cryoSPARC. Particles were automatically picked using Relion 3.0.6[ref.](*6-9*) from manually selected micrographs. After 2D classification with Relion, good particles were selected and subject to 2D classification, Ab-Initio Reconstruction and multiple cycle of heterogeneous refinement without symmetry using cryoSPARC(*10*). The good particles were selected and subjected to Local CTF Refinement with C1 symmetry, Non-uniform Refinement, resulting in the 3D reconstruction for the whole structures. To further improve the resolution, the particles of Non-uniform Refinement are subject to 3D classification and focused refinement with Relion.

The resolution was estimated with the gold-standard Fourier shell correlation 0.143 criterion(*11*) with high-resolution noise substitution(*12*). Refer to Supplemental information, fig. S1-S5 and Table S1 for details of data collection and processing.

**Model building and structure refinement**

Predicted models of F8, E4 and A22 was first obtained using alphafold2.1(*13*), which was further manually adjusted based on the cryo-EM map with coot(*14*). Each residue was manually checked with the chemical properties taken into consideration during model building. Several segments, whose corresponding densities were invisible, were not modeled. Structural refinement was performed in Phenix(*15*) with secondary structure and geometry restraints to prevent overfitting. To monitor the potential overfitting, the model was refined against one of the two independent half maps from the gold-standard 3D refinement approach. Then, the refined model was tested against the other map. Statistics associated with data collection, 3D reconstruction and model building were summarized in Table S1.

**Molecular Docking**

Docking simulation with the software Autodock Vina (Version 1.2.0) to investigate interaction of protein and ligands, according to the following procedure: (1)

Preparation of template structure of remodeled E9 and F8. We used the yeast polymerase δ structure with bound template and complementary DNA (PDB ID: 3IAY) to generate the elongation mode of F8 and E9 implemented in SWISS-MODEL (https://swissmodel.expasy.org) (*16*). The models were processed by removing existing ligands while missing polar hydrogen were added utilizing AutoDockTool software. Subsequently, saved into a dockable PDBQT format for the following virtual work. (2) Docking ligands preparation. The structure data file format of small molecules (CDVpp and AraC) were gained from the PubChem website (https://pubchem.ncbi.nlm.nih.gov) and minimized energy using MM2 algorithm. Using program optimized all above small ligands and converted files into PDBQT chemical format. (3) Set docking grid box. Imported PDBQT file of remodeled templates into the software to build up a docking grid box parameter file of E9 and F8, respectively, based on the active pocket where positive molecule (dCTP) was located. (4) Molecular docking was performed by AutodockVina and the final results were processed with PyMOL (Version 1.4.1).

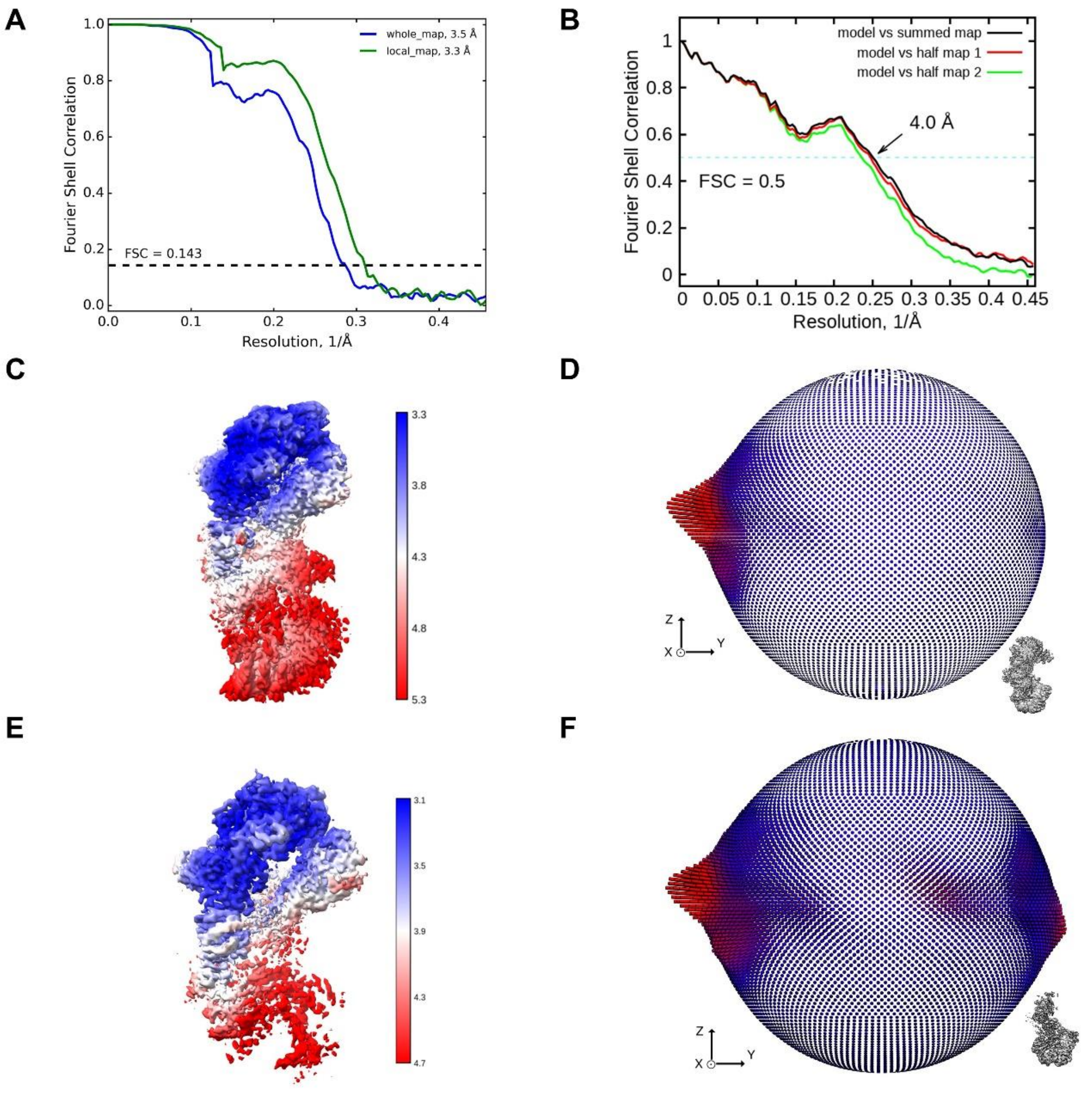


**Fig. S1**

**Cryo-EM analysis of F8/A22/E4 complex in hexameric form.**

**(A)** FSC curve. The resolution was estimated with the gold-standard Fourier shell correlation 0.143 criterion with high-resolution noise substitution. **(B)** FSC curve of the refined model versus the overall structure that it is refined against (black); of the model refined against the first half map versus the same map (red); and of the model refined against the first half map versus the second half map (green). The small difference between the red and green curves indicates that the refinement of the atomic coordinates did not suffer from overfitting. **(C)** Local resolution map for the 3D reconstruction of the overall structure. **(D)** Euler angle distribution in the final 3D reconstruction of overall map. **(E)** Local resolution map for the 3D reconstruction of the local structure. **(F)** Euler angle distribution in the final 3D reconstruction of local map.

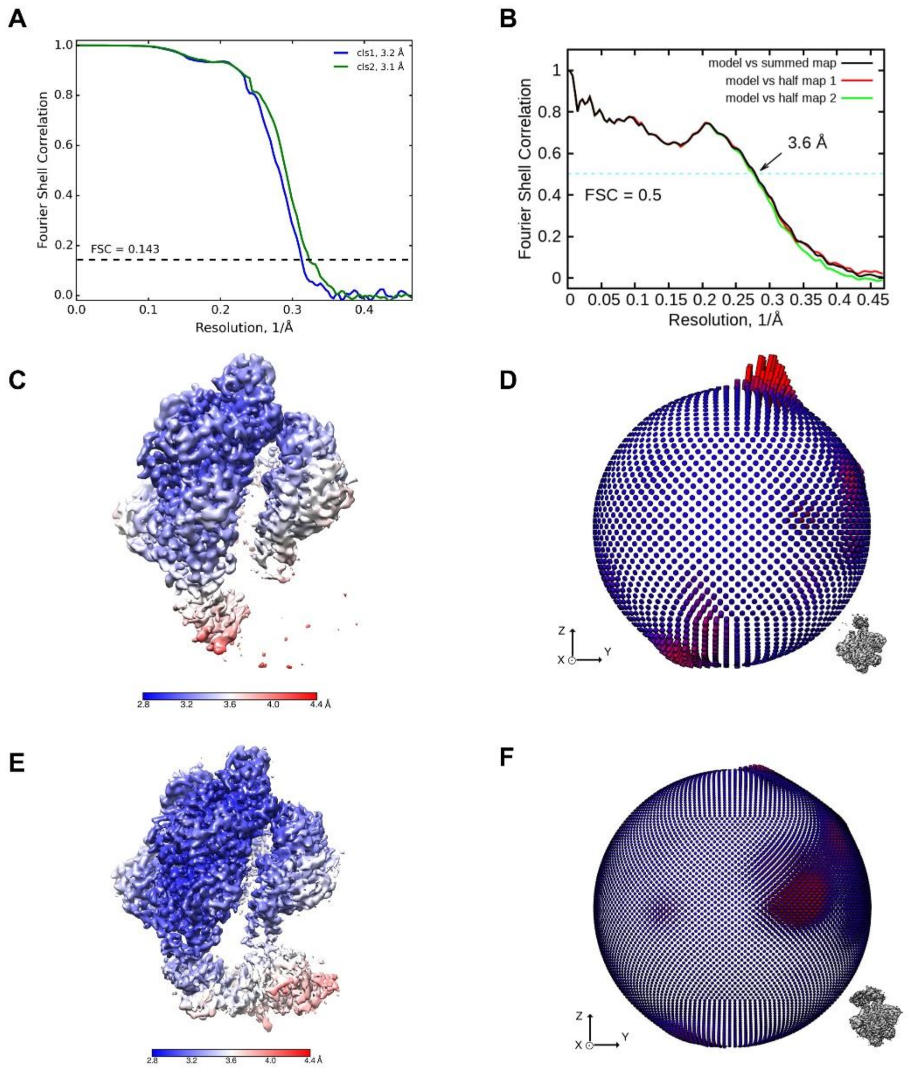


**Fig. S2**

**Cryo-EM analysis of F8/A22/E4 complex in trimeric form.**

**(A)** FSC curve. The resolution was estimated with the gold-standard Fourier shell correlation 0.143 criterion with high-resolution noise substitution. **(B)** FSC curve of the refined model versus the overall structure of class 2 that it is refined against (black); of the model refined against the first half map versus the same map (red); and of the model refined against the first half map versus the second half map (green). The small difference between the red and green curves indicates that the refinement of the atomic coordinates did not suffer from overfitting. **(C)** Local resolution map for the 3D reconstruction of the overall structure of class 1. **(D)** Euler angle distribution in the final 3D reconstruction of overall map of class 1. **(E)** Local resolution map for the 3D reconstruction of the overall structure of class 2. **(F)** Euler angle distribution in the final 3D reconstruction of overall map of class 2.

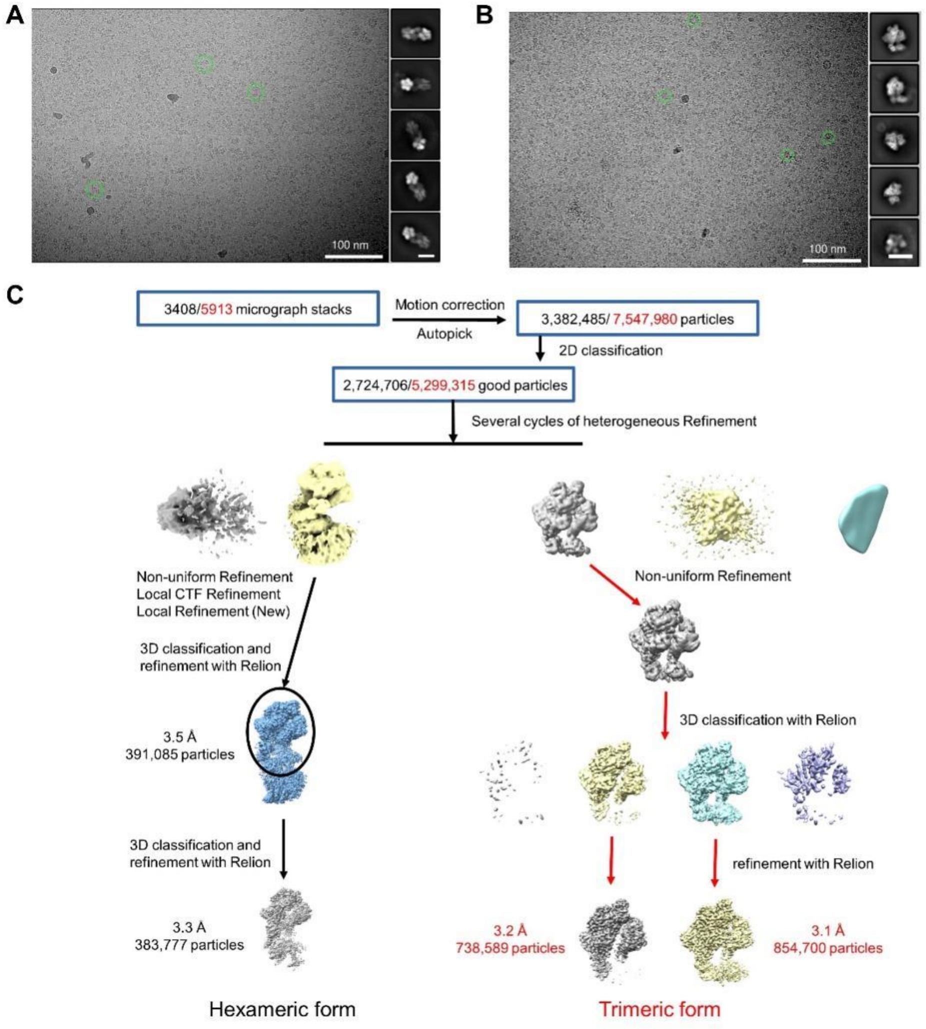


**Fig. S3**

**Flowchart for cryo-EM data processing.**

Please refer to the ‘Data Processing’ in Methods section for details.

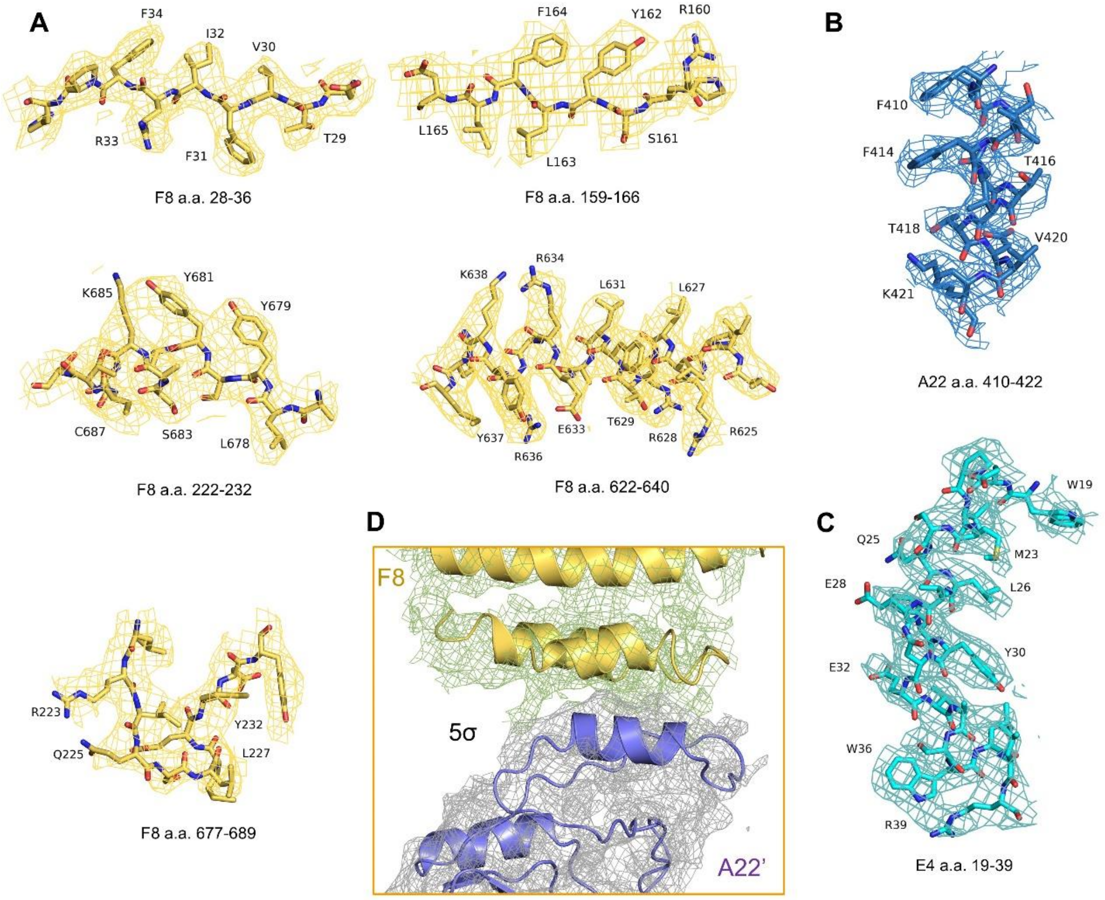


**Fig. S4**

**Representative cryo-EM density maps of F8/A22/E4 complex in hexameric form.**

**(A-C)** Cryo-EM density map of F8, A22 and E4 are shown at threshold of 7 σ. **D** Cryo-EM density map of interface between thumb domain of F8 and A22 is shown at threshold of 5 σ.

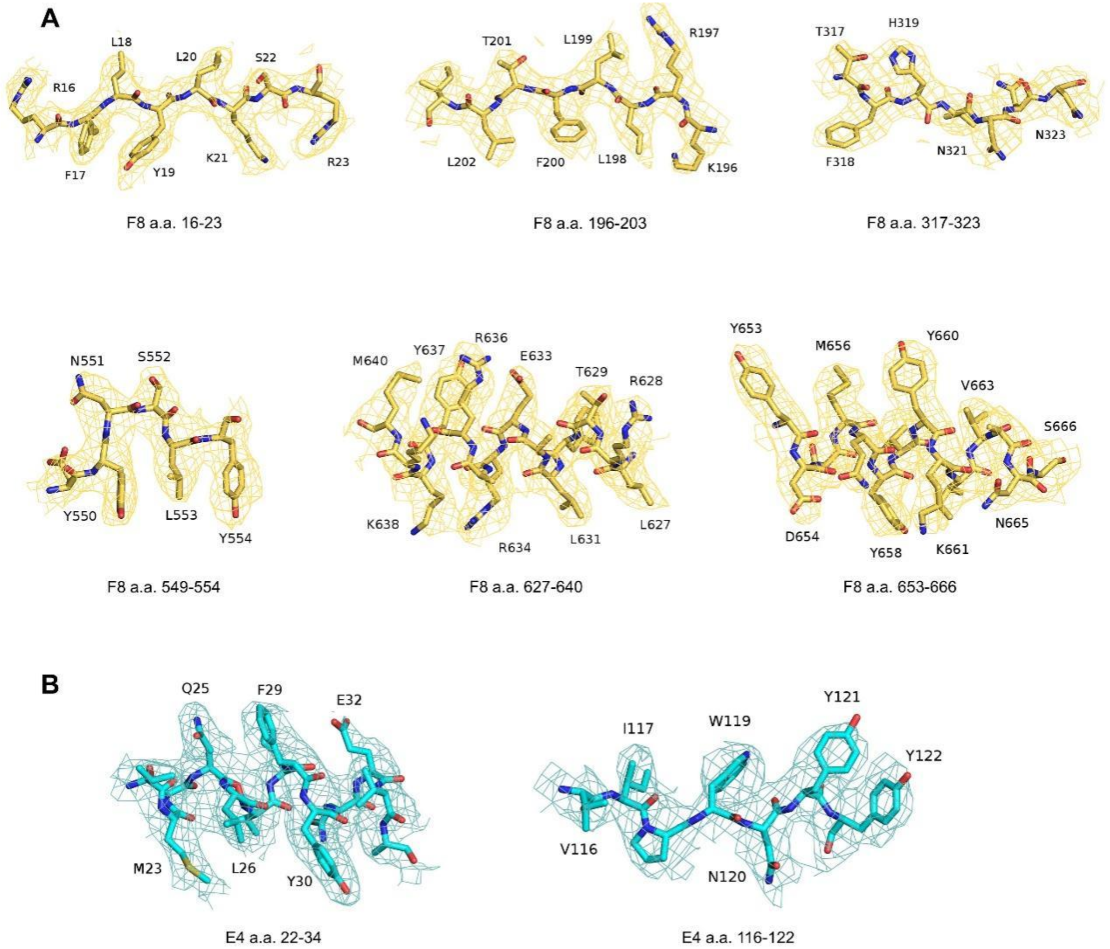


**Fig. S5**

**Representative cryo-EM density maps of F8/A22/E4 complex in trimeric form.**

**(A)** Cryo-EM density map of F8 are shown at threshold of 7 σ. **(B)** Cryo-EM density map of E4 are shown at threshold of 7 σ.

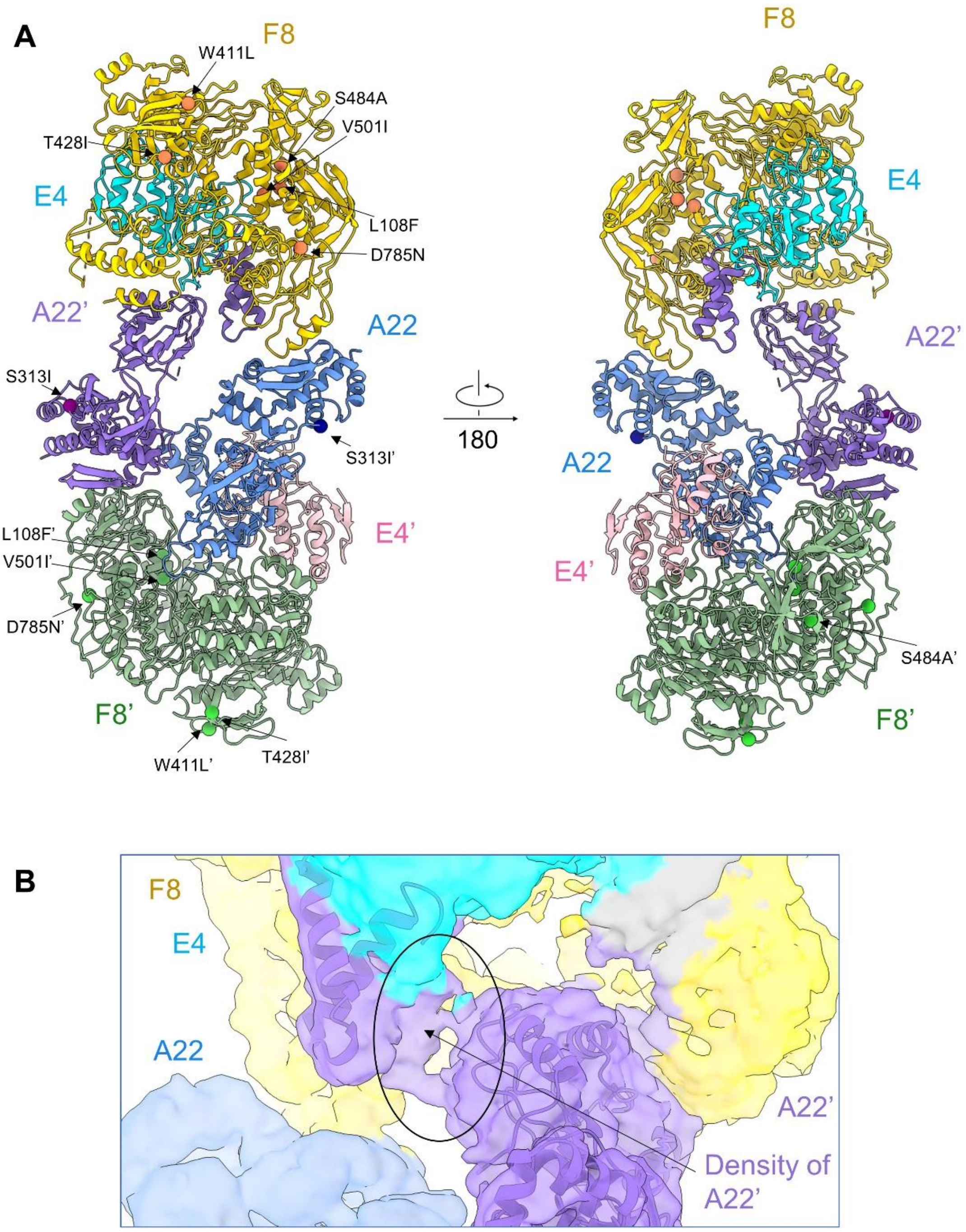


**Fig. S6**

**The distribution of mutations of F8 and A22 from 2022 West African strain sub-lineages.** (A) F8, A22 and E4 in protomer A are colored gold, blue and cyan, and the other protomer is colored green, purple and pink. Resides changed are styled sphere. Mutations on F8, F8’, A22 and A22’ are colored orange, green, blue, and purple, respectively. (B) Density of the linker between N-terminal helix and neck domain of A22'.

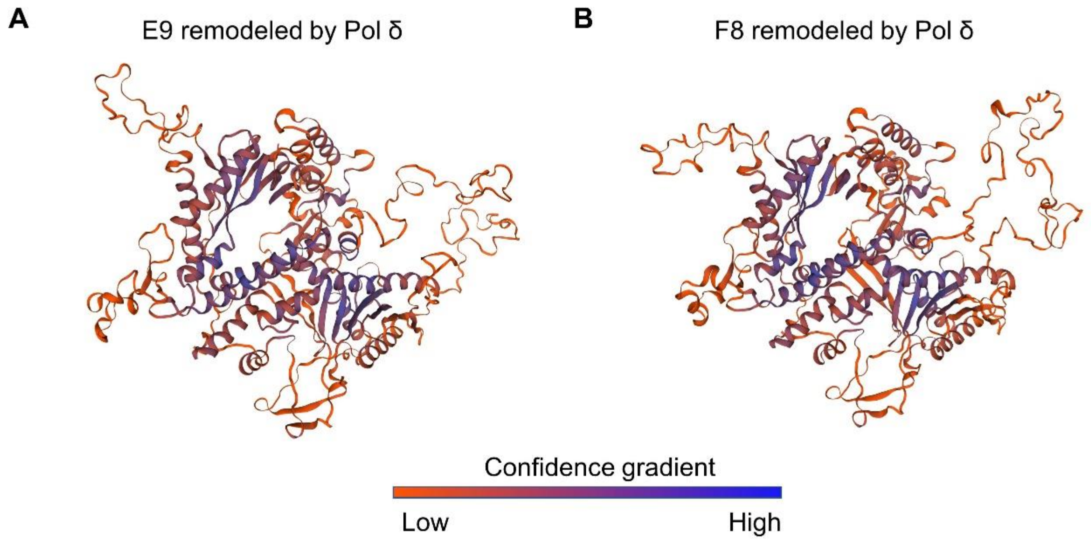


**Fig. S7**

**Close conformation of E9 in VACV and F8 in MPXV**

E9 in VACV (A) and F8 in MPXV (B) are remodeled using Pol δ (PBD ID: 3IAY) as template by SWISS-MODEL. Models are colored by confidence gradient.

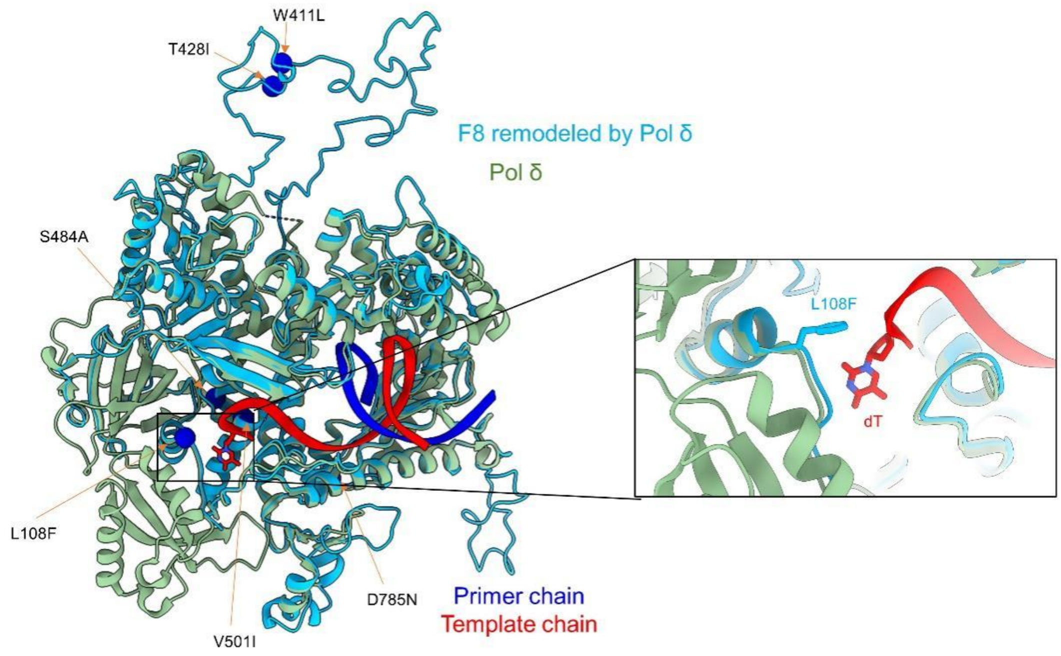


**Fig. S8**

**The distribution of mutations of F8 from 2022 West African strain sub-lineages in F8 at close conformation.**

Structural comparison between F8 at close conformation and Pol δ in complex with DNA (PBD ID: 3IAY) shows L108F may strengthen hydrophobic interaction with base of dT in DNA' template chain. F8 at close conformation and Pol δ are colored cyan and green, respectively. Primer chain and template chain of DNA are colored blue and red. Resides changed are styled sphere. F108 and dT are styled sticks.

**Table S1**

**Cryo-EM data collection, refinement and validation statistics**

| | | |
|---|---|---|
| **Data collection** | | |
| EM equipment | Titan Krios (Thermo Fisher Scientific) | |
| Voltage (kV) | 300 | |
| Detector | Gatan K3 Summit | |
| Energy filter | Gatan GIF Quantum, 20 eV slit | |
| Pixel size (Å) | 1.095 | 1.072 |
| Electron dose (e-/Å2) | 50 | |
| Defocus range (μm) | -1.4 ~ -1.8 | |
| Sample | F8-A22-E4 hexameric form | F8-A22-E4 trimeric form |
| Number of collected micrographs | 3,408 | 5,913 |
| **3D Reconstruction** | | |
| Software | Relion/cryoSPARC | |
| Sample | Overall | Overall |
| Number of used particles (Overall) | 391,085 | 854,700 |
| Resolution (Å) | 3.5 | 3.1 |
| Symmetry | C1 | |
| Map sharpening B-factor (Å2) | -90 | |
| **Refinement** | | |
| Software | Phenix | |
| Cell dimensions | | |
| a=b=c (Å) | 313.056 | |
| α=β=γ (˚) | 90 | |
| Model composition | | |
| Protein residues | 3,044 | 1,532 |
| Side chains assigned | 3,044 | 1,532 |
| R.m.s deviations | | |
| Bonds length (Å) | 0.012 | 0.019 |
| Bonds Angle (˚) | 0.960 | 0.718 |
| Ramachandran plot statistics (%) | | |
| Preferred | 83.11 | 90.05 |
| Allowed | 16.13 | 9.88 |
| Outlier | 0.77 | 0.07 |
